# RadGenNets: Deep Learning-Based Radiogenomics Model For Gene Mutation Prediction In Lung Cancer

**DOI:** 10.1101/2022.04.13.488208

**Authors:** Satvik Tripathi, Ethan Jacob Moyer, Alisha Isabelle Augustin, Alex Zavalny, Suhani Dheer, Rithvik Sukumaran, Daniel Schwartz, Brandon Gorski, Farouk Dako, Edward Kim

## Abstract

In this paper, we present our methodology that can be used for predicting gene mutation in patients with non-small cell lung cancer (NSCLC). There are three major types of gene mutations that a NSCLC patient’s gene structure can change to: epidermal growth factor receptor (EGFR), Kirsten rat sarcoma virus (KRAS), and Anaplastic lymphoma kinase (ALK). We worked with the clinical and genomics data for each patient as well CT scans. We preprocessed all of the data and then built a novel pipeline to integrate both the image and tabular data. We built a novel pipeline that used a fusion of Convolutional Neural Networks and Dense Neural Networks. Also, using a search approach, we pick an ensemble of deep learning models to classify the separate gene mutations. These models include EfficientNets, SENet, and ResNeXt WSL, among others. Our model achieved a high area under curve (AUC) score of 94% in detecting gene mutation.

## 1 Introduction

Artificial Intelligence has taken a huge leap in the field of medicine, fostering more advanced diagnostic and prognostic outcomes. We have seen AI algorithms working in a completely clinical setting to detection of Covid using binary features [1, 2, 3, 4, 5]. The future of the ‘Standard’ medical practice will be here sooner than anticipated, where a patient will be seeing a computer before seeing a doctor through advances in artificial intelligence (AI). AI systems are used extensively in medical sciences. Common applications include diagnosing patients, end-to-end drug discovery and development, improving communication between physician and patient, transcribing medical documents, such as prescriptions, and remotely treating patients.

### 1.1 Non-Small Cell Lung Cancer

Lung cancer is the 2nd most common cancer type in the world and responsible for the most cancer deaths each year, totaling 2.21 million cases and 1.80 million deaths just in 2020 [6]. Lung cancer can be further categorized into two types, small cell lung cancer (SCLC) and non-small cell lung cancer (NSCLC). Out of all lung cancer cases, NSCLC represents about 85 percent of cases [7]. The leading risk factor for causing NSCLC results from smoking and inhaling second-hand smoke [8]. In the past years, lung cancer has become more common among former smokers than current smokers. A study of more than 5000 people, diagnosed with lung cancer between 1997 and 2002 found that only 25% were current smokers, whereas more than 60% were former smokers [9]. NSCLC can be found throughout the lungs, typically on the bronchioles or alveoli. These cancerous cells eventually develop into lung nodules, small abnormal areas in the lung which can be identified with repeat CT scans of the chest [10].

### 1.2 Radiogenomics

In oncology, medical imaging is a major component in clinical decision-making. Radiogenomics focuses on correlating characteristics from medical imaging and genetic features. It can also be referred to as analyzing associations between patients’ genetics to their reaction to their radiation therapy [11]. A typical radiogenomic pipeline can involve image segmentation, feature extraction, and using statistical models to correlate the variation of clinical data [12]. The imaging features are qualitative and will assign a score to certain data parameters. Features are mathematically extracted with algorithms in radiomics. Radiomic signatures once extracted, can be utilized for several applications, including tumor categorization, survival prediction, and therapeutic response prediction [13]. The developing topic of radiogenomics includes studies that look for correlations between imaging, genomic, and molecular data [14]. Radiogenomics plays a significant role in future cancer research as it creates a new path toward generating knowledge through limited data.

### 1.3 Gene Mutation

In NSCLC, the patient’s gene structure tends to change, this change is known as “mutation”. The three major types of genes mutation are epidermal growth factor receptor (EGFR) [15], Kirsten rat sarcoma virus (KRAS [16], and Anaplastic lymphoma kinase (ALK) [17]. EGFR mutation refers to a chemical that inhibits the action of the epidermal growth factor receptor, a protein that aids in cell growth and division [18]. This mutation is found in the tyrosine kinase enzyme (From exon 18-21). Exon mutation occurs when parts of a gene (exons) are absent or deleted. If an exon mutation is found, the tumor is referred to as be EGFR mutant [19]. KRAS is a gene that produces a protein that regulates cell growth, maturation, and death via cell signaling pathways. Mutant KRAS refers to the gene’s altered or transformed form, which can enable cancer cells to proliferate even faster [20]. ALK is another gene that produces a protein that helps in cell development. Non-small cell lung cancer has been reported to contain different mutated variants of the anaplastic lymphoma kinase gene and protein. Cancer cells may grow faster as a result of these alterations [21]. The gene mutations are mutual to one another, having one over the other might affect responsiveness to targeted treatment. If either of the genes is found in a natural, unaltered form, then it is called a wild-type [22]. Therefore, due to this property of the mutation, we use the three gene mutation as our classification classes.

### 1.4 Machine Learning Approaches

Machine learning is an area of artificial intelligence (AI) and computer science that focuses on uncovering patterns in data and making future predictions using datasets and algorithms. Recent advances in deep learning have proven successful image analysis applications without the need for human feature definition [23]. Currently, deep learning methods, particularly Convolutional Neural Networks (CNN), have also been used extensively in computer vision and image processing tasks [24]. In Lung Cancer research, CNNs have mostly been investigated in the context of lung pattern categorization on CT scans[25], PET scans[26], and X-rays.

### 1.5 Models

Most of our work is done using EfficientNets (EN) [27], which have been pre-trained with the AutoAugment v0 policy [28] on the ImageNet dataset. This model family consists of eight separate models that are architecturally related to one another and that adhere to a set of scaling criteria in order to be adjusted to greater picture sizes. The smallest variant, B0, utilizes the usual input size of 224×224 pixels for its input. Larger versions, up to B7, take advantage of larger input sizes while simultaneously scaling up network width (the number of feature maps per layer) and network depth (the number of feature maps per layer) (number of layers). We use EN B0 through B6 standards. In order to include additional heterogeneity in our final ensemble, we also incorporate a SENet154 [29] and two ResNext variations that were trained with weakly supervised learning (WSL) on 940 million pictures [30] before being combined.

## 2 Related Work

### 2.1 Non-Small Cell Lung Cancer

A similar task was undertaken by researchers wanting to use the patient’s genome data in predicting recurrence of NSCLC [31]. Also using a publicly available NSCLC dataset, the researchers used a gene estimation model and another model to use the gene estimations to predict recurrence. Their maximum prediction accuracy was 83.28% using their genotype-guided radiomics method [31]. While these are very promising results, we hope to produce a model with higher accuracy through combining the genomic data with the DICOM imaging and clinical data.

Another group of researchers attempted to predict the prognosis of a patient with NSCLC by automating the evaluation of histopathology images, rather than manually evaluating them. They built an automated information pipeline to extract quantitative features(cell size, shape, distribution of pixel intensity in the cells and nuclei, etc.) from 2,186 whole-slide histopathology images. They used these features to create a model that was able to distinguish between shorter-term survivors from longer-term survivors. The digital methods used by this study were able to identify quantitative features such as Zernike shape features that may be more difficult to identify manually [32].

### 2.2 Radiogenomics

Plodkowski et al found evidence to support using radiogenomics to prove the presence of NSCLC in patients [33]. In this paper, researchers analyzed RET and ROS1 rearrangements against an EGFR mutant group in patients with lung adenocarcinoma. In conjunction with their genomic sequences, the researchers also utilized CT imaging of the patients’ lungs. This radiogenomic technique allowed the researchers to find a statistically significant difference (p = 0.04 at 95 percent confidence) in peripheral tumors appearing in CT scans when patients had the ROS1 rearrangement present in comparison to the RET and EGFR groups. Similar radiogenomic techniques as these researchers can be used to find a stronger connection between genomic sequencing and CT imaging results to predict survival risk and reoccurrence of tumors.

Radiogenomics has also been applied to modeling risk and dangers in other individuals suffering from cancer. In a study by Zhao et al, they analyzed patients that suffered from prostate cancer and their risk of receiving radiotherapy toxicity [34]. Utilizing a normal tissue complication probability (NTCP) model, they compared patients to find trends between single nucleotide polymorphisms(SNP) and the increased probability of developing radiotherapy toxicity. Kerns et al found a positive correlation between the number of SNP occurrences and the risk of radiotherapy toxicity [35]. Radiogenomics was further used alongside NCTP to find how tumors in these patients respond to radiotherapy given the number of SNP occurrences, being less effective as the number goes up as well [36]. Radiogenomic techniques in these previous papers can be used to identify trends between recurrence of NSCLC and genomic sequences linked with this risk of recurrence.

### 2.3 Machine Learning Approaches

In a related study, deep learning networks were used to analyze time-series CT scans of patients with locally advanced non-small-cell lung cancer to predict clinical outcomes. The study consisted of two datasets (dataset A and dataset B), comprising in total of 268 with stage III NSCLC for the analysis. They developed a model utilizing convolutional and recurrent neural networks (CNN and RNN), using a single seed-point tumor localization to accurately predict the survival rate and other cancer-related outcomes such as progression, recurrence and metastases [37].

Another study utilized a deep convolutional neural network to detect the two most prevalent subtypes of lung cancer: adenocarcinoma (LUAD) and squamous cell carcinoma (LUSC). The study used 1,634 whole-slide images which consisted of 1,176 tumor tissues and 459 normal tissues. The model was trained by using non overlapping 512 × 512 pixel tiles. They also trained the model to predict mutated genes in LUAD and found that six out of the top ten commonly mutated genes can be predicted from pathology images [38].

Moreover, another study attempted to predict cell proliferation in cases of non-small cell lung cancer. The model used a random forest feature selection algorithm (RFFS) to reduce features amongst 245 CT scans. The study used the Ki-67 proliferation index which measures how fast cells grow. After using six machine learning models, 103 radiomics features were extracted along with 20 optimal features. The random forest-based radiomics classifiers beat other combinations of classifiers in predicting Ki-67 expression levels [39].

## 3 Methodology

### 3.1 Dataset

The [Dataset](https://wiki.cancerimagingarchive.net/display/Public/NSCLC+Radiogenomics) was obtained from The Cancer Imaging Archive, and was downloaded using the NBIA Data Retriever [40]. The Dataset’s main purpose is to discover the underlying relationship between genomic and medical image features, as well as the development and evaluation of prognostic medical image biomarkers. It’s compromised of three main parts: The imaging data, the RNA sequencing, and gene mutation data, and the clinical data.

The imaging data is made up of Computed Tomography (CT), Positron Emission Tomography (PET)/CT images, semantic annotations of the tumors as observed on the medical images, and segmentation maps of tumors in the CT scans, and quantitative values obtained from the PET/CT scans. The sequencing and mutation data are collected from samples of surgically excised tumor tissue. The clinical data includes much information on each patient as well as their recurrence outcomes.

In selecting which features of the clinical data would be used for the Machine Learning models, a correlation matrix was used to determine which features were most correlated with Recurrence.

### 3.2 Pre-processing

The raw clinical data contained many missing values and had to be pre-processed accordingly. Missing values for the continuous variables such as Weight and Pack Years were replaced with the mean, as the variables had a fairly normal distribution. Missing values for the categorical variables were replaced with “None”, as the quantity of data available prevented a replacement of the existing values. The Recurrence and Death dates were binned into categories of 5-year increments over a decade, as the limited quantity of our data prevented us from making accurate predictions of the exact date when a patient will die of lung cancer will reoccur. The continuous features of the data were then normalized, and the categorical features were either ordinally or one-hot encoded. The Smoking status feature was also split into three separate columns of former, non, and current smokers to compare their correlation with Recurrence individually. The correlation of the clinical data is shown in Figure 1.

**Figure 1:**
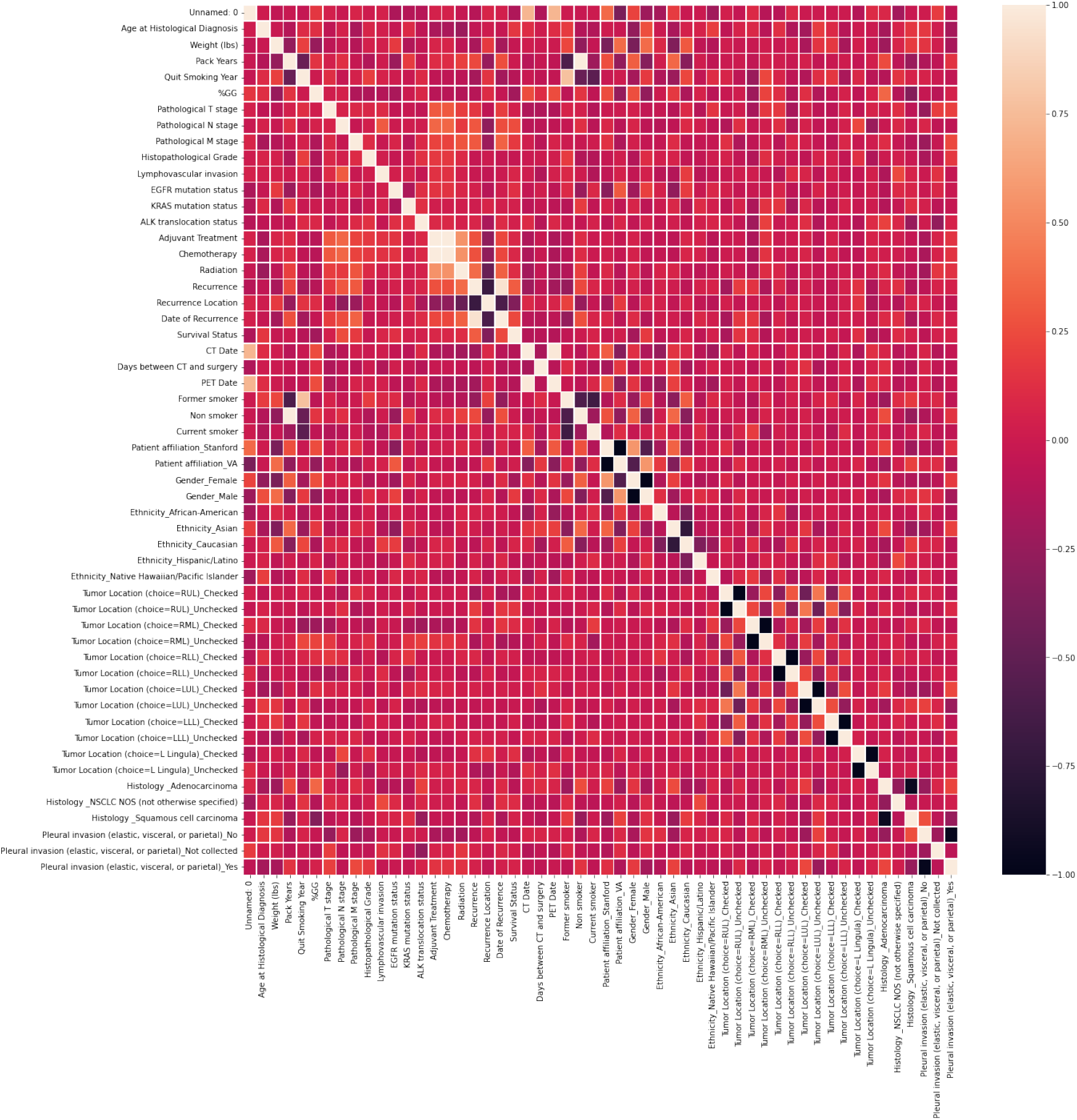
Clinical data correlation heat map.

The genomic data were pre-processed by replacing all N/A values with zero, dropping invariant features, and normalizing each feature across rows. Then, the 100 most variable features were selected. The RNA sequencing data also only had recorded genomes for 130 of the patients, which would then cut the clinical data down roughly by half. The correlation heat map between the genes and the patients is shown in the Figure 2.

**Figure 2:**
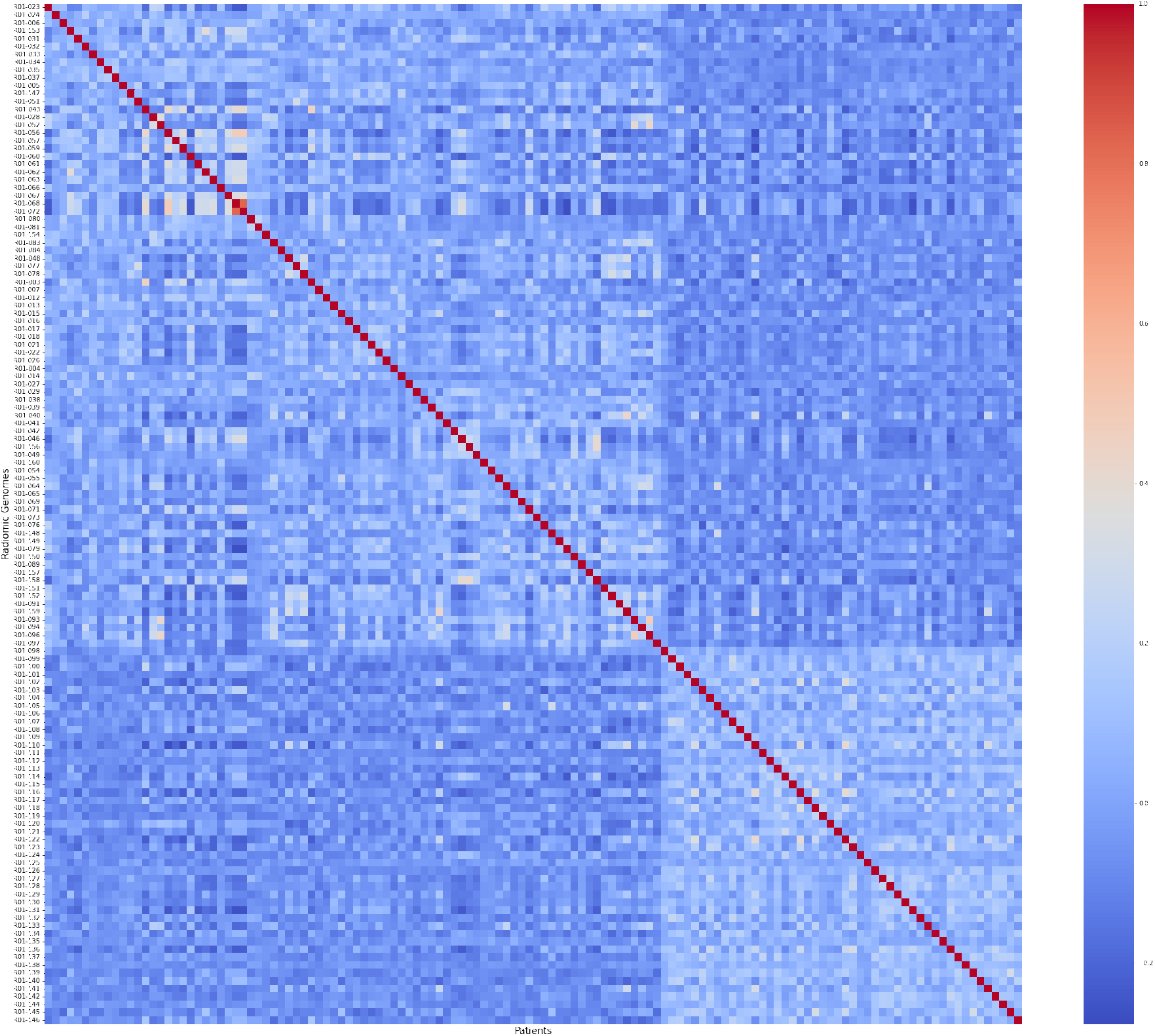
Genomics features.

The lung cancer scans were in digital imaging and communications in medicine (DICOM) format, which is the standard for medical images. The images’ pixel values were calibrated to Hounsfield Units (HU), a measure of radiodensity and the general unit of measurement for CT scans. The conversion was performed by implementing a linear scaling step on each image of the CT scan and this allowed filtering of any irrelevant substances present in the images. Because the CT scans had a variety of resolutions, this can cause an issue in the model training process. To avoid this, all the images were resampled to have a set number of 256 slices, which will be eliminating a source of error and ensuring consistency amongst the data.

### 3.3 RadGenNet Pipeline

Our pipeline is categorized into two parts, as shown in the Figure 3. For part 1, we develop a series of convoluted neural networks for the classification of the epidermal growth factor receptor (EGFR), Kirsten rat sarcoma virus (KRAS), and anaplastic lymphoma kinase (ALK) gene mutation cases. In part 2, we build a dense neural network for the clinical data and genomics data.

**Figure 3:**
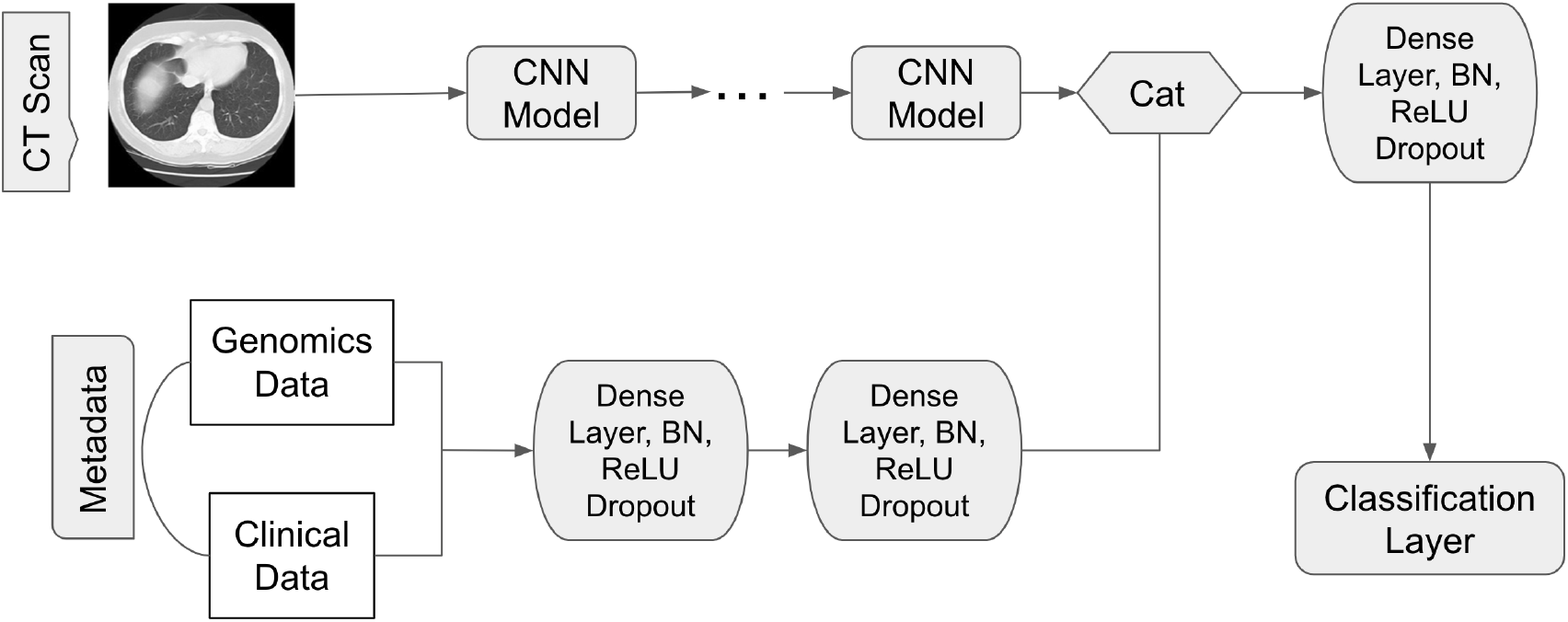
Our approach for combining CT Scan medical image data and meta data.

Initially, in the first step, we train our CNNs on the pre-processed DICOM scans (part 1). Then we hold on to the CNN weights and link the clinical and radiomics data-dense neural network on top of the CNN weights. Then in the second part, we only run the clinical and radiomics data-dense neural network and final classification layer.

### 3.4 CNN Training

We used Adam optimization to train all of our models for a total of 100 epochs. We used a weighted cross-entropy loss function, in which underrepresented classes are more often seen in the training set due to their greater weighted frequency. In each class, the total number of training pictures is multiplied by a factor where N is the number of scans in each class in class, we found it best to deal with the severity of balancing. We discovered that it was the most effective. With the same balancing weights, we also attempted to apply the focused loss [41] method but found no increase in performance. The batch size and learning rate are chosen in accordance with the GPU memory limitations of the respective architecture. Every 25 epochs, we cut the learning in half. Every ten epochs, we assess the model and preserve the one with the best mean sensitivity (or sensitivity to noise) (best). In addition, for every 100 epochs of training, we preserve the most recent model (last).

### 3.5 Meta Data Architecture

For part 2, the preprocessed clinical and genomics data is loaded into a two-layer neural network with 256 neurons on each layer, which is then trained on the data. The batch normalization, ReLU activation, and dropout with p = 0.4 are all included in each layer. Following global average pooling, the output of the network is concatenated with the feature vector of the CNN. After that, we apply a second layer that includes batch normalization, ReLU, and dropout. As a starting point, we employ 1024 neurons, which are scaled up for bigger models by following the scaling guidelines for network width provided by EfficientNet. The classification layer is then added at the last of it.

### 3.6 Training and Ensembling

During part 2, the weights of the CNN are kept the same. Metadata layers, such as the two-layer network, layer following concatenation, and classification layer, are trained for 50 epochs with a learning rate of 0.00001 and a batch size of 20.

Last but not least, we assemble a big ensemble comprising all of our previously trained models. We use an approach in which we pick the best subset of models based on their performance in cross-validation tests [42].

### 3.7 Method Validation

We analyze the mean sensitivity for training with just input scans and the mean sensitivity S for training with extra metadata for assessment. Table 1 shows the results of cross-validation with individual models as well as our ensemble of models. In general, bigger ENs outperform their smaller counterparts. The process of assembling results in a significant improvement in performance. Our ideal ensembling method results in a marginal improvement in performance.

**Table 1:**
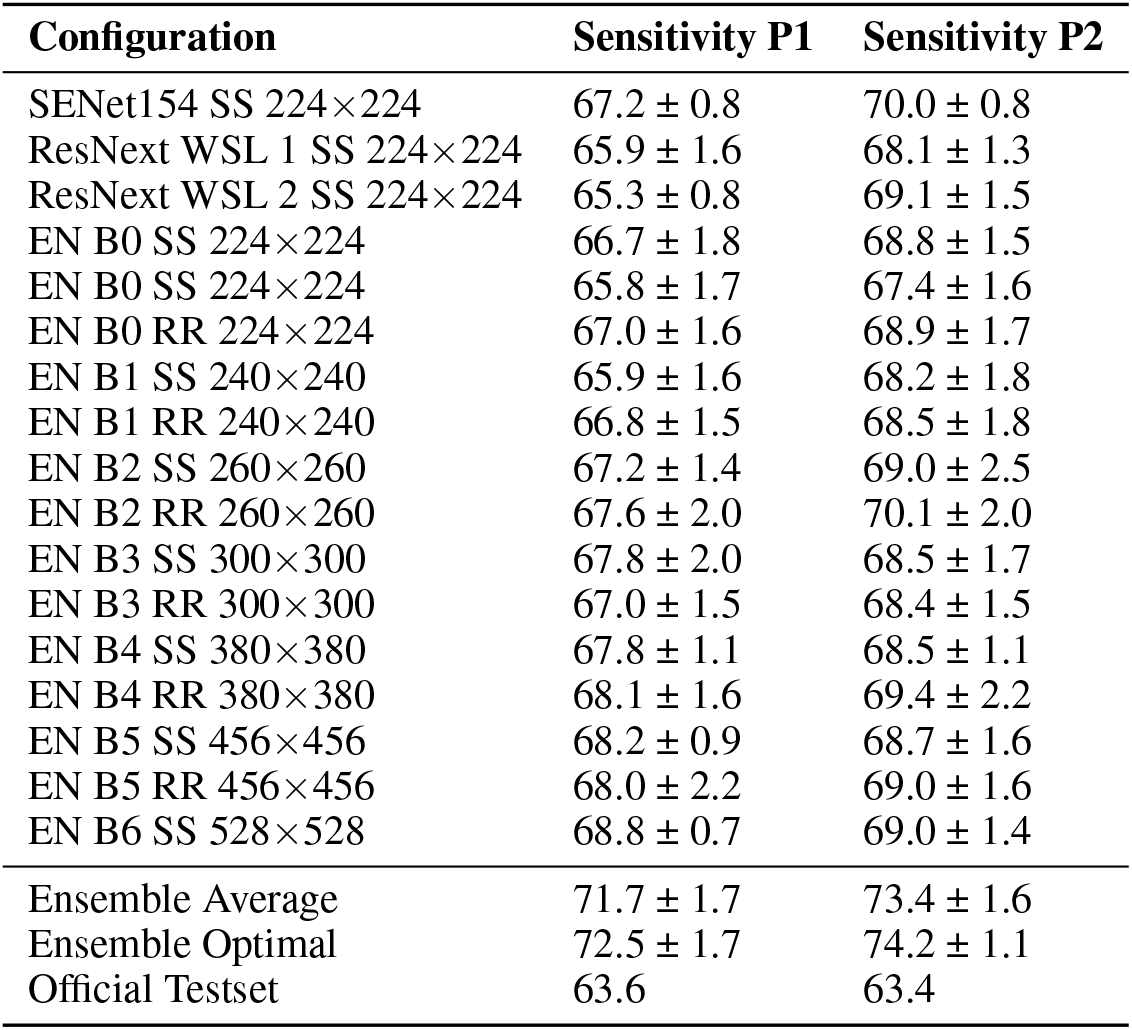
All cross-validation results for various models are included. The values are expressed as the mean and standard deviation across all five CV folds, expressed in percent. The term “ensemble average” refers to averaging across all predictions from all models in a given situation. The term “ensemble optimal” refers to averaging across the models that we discovered when searching for the optimal subset of configurations using our search approach. P1 refers to Part1 without metadata, while P2 refers to Part 2 with metadata.

## 4 Results

Our methodology consisted of two Deep Learning models getting fussed together, so it was important to see how each model is performing independently and in what ways does it affects the pipeline when combined. In Figure 4, we have shown the metrics of our Metadata Deep Neural Networks. We used the Jaccard similarity coefficient score and loss function to evaluate the model. We achieved an train accuracy of 96% and validation accuracy of 95%, thus, the performance on the test set is slightly lower than the performance obtained from cross-validation. Using the ensemble with the best model checkpoints, we were able to get some excellent performance results on the test set. Table 2 provides multiple metrics for the performance of our model with and without the metadata Dense Layer included. We take into consideration the AUC, the AUC with a sensitivity higher than 80% (AUC-S), the sensitivity, and the specificity of the test results. However, it should be noted that the sensitivity provided here is calculated differently than S.The performance of part 1 is much lower than the performance of part 2, which provides us the confidence in our multi-modal pipeline methodology.

**Table 2:**
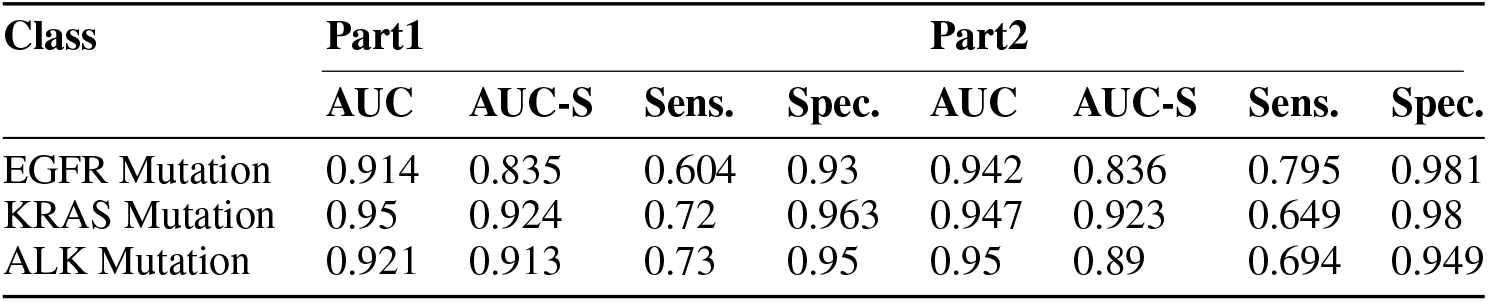
The results of the test set for each class are shown below. We take into account the AUC, the AUC for a sensitivity greater than 80% (AUC-S), the sensitivity, and the specificity of the test. It should be noted that the sensitivity offered here is derived in a different way than S. Percentages are used to represent the values.

| Class | Part1 |  |  |  | Part2 |  |  |  |
| --- | --- | --- | --- | --- | --- | --- | --- | --- |
|  | AUC | AUC-S | Sens. | Spec. | AUC | AUC-S | Sens. | Spec. |
| EGFR Mutation | 0.914 | 0.835 | 0.604 | 0.93 | 0.942 | 0.836 | 0.795 | 0.981 |
| KRAS Mutation | 0.95 | 0.924 | 0.72 | 0.963 | 0.947 | 0.923 | 0.649 | 0.98 |
| ALK Mutation | 0.921 | 0.913 | 0.73 | 0.95 | 0.95 | 0.89 | 0.694 | 0.949 |

**Figure 4:**
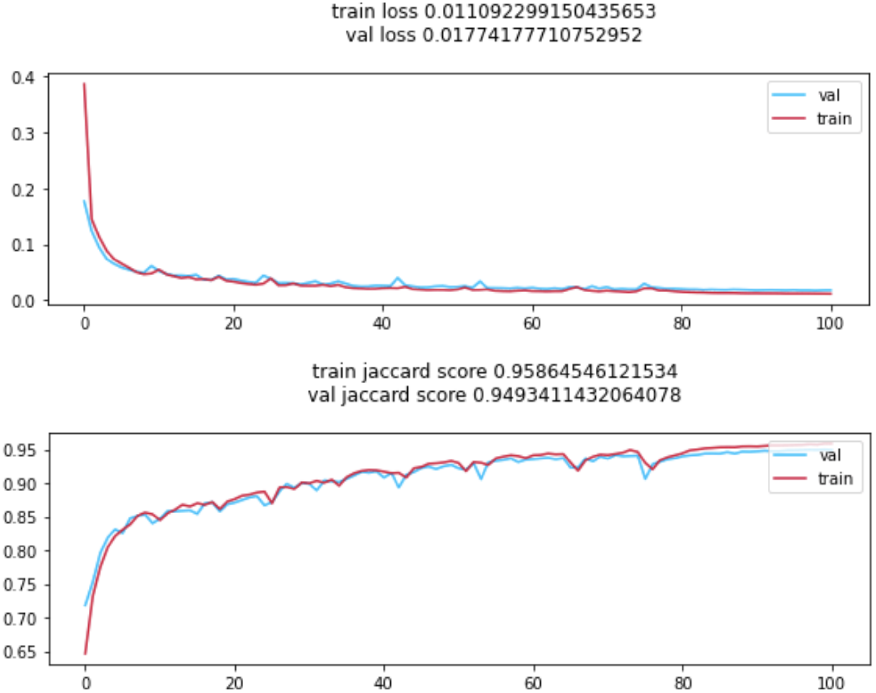
Results from the Metadata deep layer neural network architecture.

## 5 Ethics and Bias

Since a similar type of data sample is being collected, that too from an external source, as a result, “learnt” machine learning models are not generalizable. It is unconfirmed whether or not the patients had complete authorization over their records being made public which could indicate a PHI leakage. One of the biggest challenges yet remains the generalizabilty of our AI model [43]. In other words, the machine’s predictions aren’t as accurate for specific groups, and they may conflict with medical professionals’ advice if the data used to train these models isn’t annotated [44].

With whatever results are found, it’s also important to recognize a possible gender and/or ethnicity bias and how it might have affected the results. The objectivity of a supervised ML model is limited by the objectivity of the training data. Bias may have been introduced due to the characteristics of the population from which the data was collected or prejudice of the expert annotator involved [45]. 64% of the patients were male and the other 36% were female which was perceived to be balanced. Therefore, even with a thorough analysis of the algorithms and data sets, it may be difficult to eliminate every undesirable bias, especially when AI systems learn from data sets, which contain previous biases.

## 6 Conclusion

We took three different types of data and used them all together to make our predictions. Overall, we find that EfficientNets performs well for the prediction of gene mutation in lung cancer. Various EfficientNets were featured in our final ensembling method, with the larger ones outperforming the others in terms of performance. This suggests that a variety of input resolutions is a desirable option for gene mutation classification over huge data since it may cover a wide range of scales. In addition, the SENet154 and ResNext models were automatically picked for the final ensemble, indicating that a degree of heterogeneity in terms of architecture is beneficial to the experiment.

## 7 Acknowledgement

We would like to acknowledge the Drexel Society of Artificial Intelligence for its contributions and support for this research.

